# The practical value of an explicit response model in quantitative uses Biacore™ biosensors

**DOI:** 10.1101/294843

**Authors:** Jason M. Whyte

**Affiliations:** ARC Centre of Excellence for Mathematical and Statistical Frontiers, School of Mathematics and Statistics, University of Melbourne 3010, Victoria, Australia; Centre of Excellence for Biosecurity Risk Analysis, School of BioSciences, University of Melbourne 3010, Victoria, Australia

**Keywords:** Biacore™, surface plasmon resonance biosensor, biomolecular interaction models, kinetic experiments, chemical kinetics, indirect measurement, model structure specification

## Abstract

Suppose we aim to use data obtained by studying a biomolecular interaction system with a surface plasmon resonance (SPR) biosensor in quantifying some system feature. We assume a parametric mathematical model for biosensor response due to sources of mass such as analyte-ligand complexes. Some parameters represent interaction features, such as rate constants. Whenever we attempt to estimate parameters from data, we may obtain multiple estimates, regardless of the amount and quality of data. Inconveniently, we may be unable to distinguish between alternatives. This is problematic when alternative parameter values lead to very different predictions of system behaviour for a situation where we lack data. Anticipating this issue prior to data collection allows us to redesign the combination of planned experiments and model, replacing a certain failure to achieve our study’s aim with the possibility of success. The literature on SPR biosensors (and computational biology more generally) has paid little attention to this matter. In order to remedy this, it is appropriate to begin with a consideration of the assumed models. These are rarely specified completely, causing ambiguity that impedes scrutiny of their properties and comparison with other models. We demonstrate this by reviewing some model types seen in the Biacore™ biosensor literature. We propose to eliminate model ambiguity by providing a suitable framework for specifing models for biosensor data. This framework will aid future efforts to compose models for data arising from particular interaction mechanisms in a form that is amenable to scrutiny. We expect that the issues raised here will have relevance to the modelling of data obtained from other apparatus employed in quantifying binding behaviour.

## 1 Introduction

Biosensors are established as a standard means of monitoring the interactions of biomolecular species. The Biacore™ marque of “flow-cell optical biosensor”,^1^ released in 1990, was the first commercial offering. Units in this line did not require labelled interactants, giving them certain advantages over those apparatus which required labels ([2]). The rapid uptake of various Biacore™ units has led to their widespread and continuing use in published studies over a range of applications in molecular biology.^2^

One use of Biacore™ data is in the quantification of features of some biomolecular interaction system, such as through estimation of its rate constants. This process requires a parametric model for the data. The problem of estimating parameters from data is generally referred to as an ‘inverse problem’.

The biosensor literature relating to inverse problems (and the computational biology literature more generally) has given much more consideration to data than models. Some authors (for example, [3-5]) have provided guidance on experimental practice and suitable experimental conditions to improve data quality. This advice may be aimed specifically at increasing the accuracy of parameter estimates ([6]).

However, in any inverse problem, features of the assumed model may ensure that parameter estimation cannot return a unique estimate for each parameter. This can occur regardless of the amount and quality of data. Non-uniqueness of parameter estimates is problematic when alternative estimates lead to very different predictions of system behaviour for a situation where we lack data. If distinguishing between equally-likely alternative estimates to ascertain the ‘true’ parameter vector is impossible, the result is an inconclusive study. Specifically, we cannot credibly use our model in predicting system behaviour.

We may be able to anticipate this problematic outcome prior to data collection through inspection of the assumed model. Suppose we test a model for the property of (global a *priori*) identifiability (see Section 2.8). If the result is negative, then it is almost certain that parameter estimation cannot return a unique estimate for each parameter. Further, the test can foresee the event where it is almost certain that there will be a continuum of equally-valid estimates for at least one parameter. In this case we will be unable to discern the true parameter value, and accordingly, a planned study will not deliver the desired information.

To the best of our knowledge, there are few considerations of identifiability as it pertains to models for data from biosensor experiments. The earliest published efforts ([7-9]) focused on formulating testing methodologies suitable for a class of mathematical models appropriate for modelling certain experimental situations. Any such testing method was hitherto absent from the systems theory literature. The particular focus of these works was on ‘kinetic’ experiments used to collect data for the estimation of interaction rate constants.

The most developed version of the testing theory appeared in [10]. This thesis also demonstrated that alternative forms of a model for data from kinetic experiments can have different properties. In particular, the work considered models for data obtained from the ‘simple bimolecular’ interaction mechanism.^3^ It showed that a four-parameter model lacked the property of identifiability, yet a three-parameter form derived from it was identifiable.

By its nature, the testing of models for identifiability is a technical — often laborious — matter. However, we may demonstrate a case where a particular model lacks the property of identifiability with a web application [11]. In this ‘app’ the user can experiment with a four-parameter form of a model for the simple bimolecular interaction which produces simulated data. The app provides two sets of input fields for parameter vectors, Inputs 1 and 2. These inputs are associated with simulated time courses of response, Outputs 1 and 2 respectively, plotted on the same axes. After Input 1 is entered, the app provides a distinct vector that will reproduce Output 1. The user can confirm this by setting Input 2 to the supplied values, noting that it causes the superimposition of Output 2 on Output 1. The user may repeat this process for various values of Input 1. The wide range of distinct values produced by the app is a consequence of the existence of infinitely-many parameter values that produce the same model output.^4^

Whilst simulated data can suggest that a given model lacks identifiability, it is not a formal test for the property. However, in order to conduct an *a priori* identifiability analysis of a model for SPR biosensor data, we may first have to expend some effort to understand the features of our assumed model. Otherwise, the practical meaning of the results may be unclear. This opacity follows from uncertainty over the information contained within an assumed model. Rarely does the literature specify a model completely. Such underspecification makes it difficult to interpret a model in those cases where a paper suggests (or omits) details of the dynamics of the interaction, or features of experiments and associated biosensor response, rather than making them explicit.

We can illustrate the variation in detail associated with models through a survey of the literature. Frequently, papers only provide the name of the interaction mechanism fit to data (such as [12, 13]). (We will cite more examples of this tendency shortly.) Some papers also include the chemical equations associated with interactions; see, for example, [14, 15]. At times a work will show a model for the interaction kinetics, and omit an expression for biosensor response. The model may appear in the form seen in the BIAevaluation software that accompanies Biacore™ units ([16]), or some other form ([17]). Alternatively, a paper may present a system of ordinary differential equations (ODEs) for response. This system may relate to the association phase of a kinetic experiment ([18]), and less commonly will also model the dissociation phase response ([19]). (We will explain the phases of kinetic experiments in Section 2.3.) Other papers provide a response function that is an explicit function of time ([20]).

Many of the alternative model forms are not easy to interpret. To appreciate this, consider consider the nature of the biosensor response. It has multiple components due to various sources of mass in an experiment. Some ‘uninformative’ components do not relate to the progress of interactions. However, when models in the literature include some expression for response, they often do not articulate which components are included, or how they are composed. This lack of detail may obscure which species make a contribution to modelled response, and the manner in which they do this. It is advantageous to have a model that we can clearly relate to our data, as we will demonstrate shortly.

It may appear that certain situations do not require a completely specified model. Consider the case where one intends to use BIAevaluation software in fitting a model to Biacore™ data. The user may select an interaction mechanism from a menu. Various recent protocols ([21-23]) or research papers ([24, 25]) consider this approach to model fitting. These works restrict their attention to two relatively simple and commonly used interaction models (outlined in Section 2.2), presenting only their names. However, each of these references provides direction on how to address a preliminary question:

(Q1) How should we process response to obtain a time course of data that is appropriate for our assumed model?

so that parameter estimation can follow.^5^

The approach to data processing outlined in the works cited above is not appropriate for every situation. For example, it is inappropriate for a case where immobilised ligand dissociates from the sensor surface.^6^

We see that the literature, by focussing on the more common interaction models, does not address Q1 in a general manner that can inform the use of other models. While a suitably experienced experimentalist will employ expert knowledge to answer Q1, the less-experienced user might not reach an answer so readily. Further, the manner in which models are presented in the literature often does not assist us in answering Q1. This is especially the case when we consider an interaction model that is more complex than the standard models. Later we will show that a fully specified model can guide the user towards appropriately answering the question.

A theoretician may find that an incompletely specified model raises a variety of other questions. The answers to these questions may be of considerable practical importance. Such questions include:

(Q2) What do the model terms and parameters represent, and what units do they have?

(Q3) Is this model related to another model that employs a different set of mathematical relationships, or which uses alternative terminology?

(Q4) Can we simplify the model to one of fewer terms or parameters, making it possible to fit the model more quickly to suitably processed data?

(Q5) In the interests of evaluating our experimental design, how should we test the model for particular properties, such as global *a priori* identifiability?

(Q6) To which experimental situations are our theoretical insights from Q5 applicable?

(Q7) Do alternative forms of the model have differing properties, in particular, as these properties pertain to parameter estimation?

We show the usefulness of these questions in a simple case in Figure 1. There may be additional connections between the questions. For example, the answer to Q5 may encourage us to reconsider Q4.

**Fig. 1.**
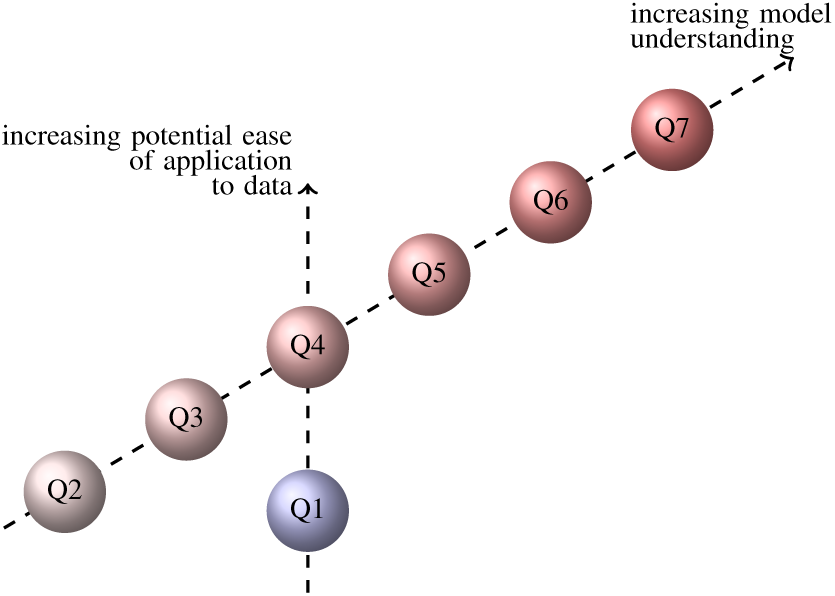
Graphical depiction of how the answers to questions Q1-Q7 may improve our ability to understand and appropriately use a model for biosensor response. In practice, if the answer to a question causes us to modify a model, this information may encourage reconsideration of other questions.

A completely specified model will make the answers to Q1 and Q2 obvious, and aid us in considering Q3-Q7. Such a model is also necessary should one not have dedicated software for fitting models to data. Even if such software is available, suppose that it does not include a model for some novel interaction mechanism or experimental situation. In such a case we must supply a model in sufficient detail to enable its addition to the software.^7^ One example of where this is necessary arises in the study of multivalent interactions:

> “Unfortunately, most models provided for SPR data analysis assume 1:1 or at most 2:1 (ligand:analyte) binding.” ([19]).

At present, completely specifying certain models is challenging. The literature does not provide specific instruction on how to specify a model. In the absence of such guidance, the literature employs a variety of alternative formalisms and terminologies. (This observation prompted Q3.) Further, to the best of our knowledge, no previous work has attempted to reconcile these alternatives. Here we provide a framework for resolving this matter. We propose a general method for specifying models of biosensor response in a transparent manner. By combining this template with the particulars of some specific interaction mechanism, one will produce a fully specified interaction model. One may then proceed to address Q1-Q7.

This paper is organised as follows. In Section 2 we present features of Biacore™ biosensors, matters relating to data collection, and models for data in more detail. Through this we formalise certain assertions regarding biosensor response and show when they are equivalent.^8^ We have particular use for a not-so-commonly cited relationship between response due to a species and molar concentration of that species from the literature.^9^ We employ this relationship in Section 3 where we propose a general means of relating response to the dynamics of some assumed interaction mechanism. We will see later that our approach is more transparent than others. Further, we demonstrate the usefulness of this representation with an example that shows how we may profitably simplify an initial model. We discuss the value of our approach and comment on future uses in Section 4.

We expect that the general thrust of our arguments will also have relevance to other biosensors for which models of response are not clearly specified. This is particularly likely when the biosensor has multi-component response (as we will see in Section 2.4) as this presents a particular opportunity for model ambiguity.

We conclude this section by introducing notation we will employ in defining models subsequently.

### 1.1 Notation

Let 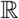 represent the set of real numbers and 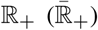 be the subset of 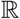 containing only positive (non-negative) reals. Let 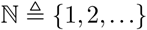 represent the natural numbers. For some 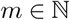, let 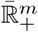 represent the set of vectors of *m* components, where each component belongs to 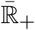. For some 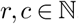, let 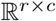 represent the set of matrices having *r* rows and *c* columns where each element belongs to 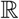. We will use a bold symbol in lower case (such as x) or upper case (such as A) to denote a vector or matrix respectively. We use A^T^ to denote the transpose of matrix A. If x is a column (row) vector, then x^T^ represents x written as a row (column) vector.

As we will make general arguments about types of chemical species later, here we define notation for their properties. Let us consider some species X, having molar mass *M*_*X*_ (g *⋅* mol^−1^). When the total mass of X is *m*_*X*_ (g), we write the number of moles of X as *N*_*X*_ (mol), where

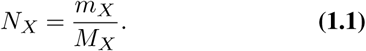

If we have *N*_*X*_ moles of X in some volume *V* (dm^3^), we represent the molar concentration of X by [X] (mol ⋅ dm^−3^), defined by

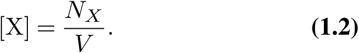

We define a volume mass concentration *c*_*X*_ (g. dm ^−3^) for the mass of X in volume *V* by

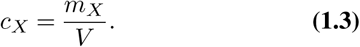

## 2 Features of Biacore™ biosensors and modelling biosensor response

We begin this section with an overview of Biacore™ biosensors and present two relatively simple and widely cited descriptions of biomolecular interactions to give the flavour of the area. We proceed to outline kinetic experiments, catalogue the components of biosensor response, and formulate a general response equation. These preliminaries inform a discussion of how we may process response data to remove uninformative components.

To demonstrate the diversity of aspects of the literature, we present some relationships between response due to a species and the amount of that species, and show how these are related. We draw on these relationships in reviewing types of models for response from the literature in order to demonstrate their variability and limitations. We conclude the section with an overview of the testing of models of Biacore™ data for the property of global *a priori* identifiability. This task was the primary motivation for this investigation of model representations, and such testing will be aided by the results of this study.

### 2.1 An overview of the apparatus

Biacore™ biosensors monitor the interactions of at least one species bound to a sensor surface (‘immobilised ligand’) with some species present in a solution made to flow over the surface (‘analyte’). Contact between a sensor chip and an integrated fluidics cartridge (IFC) that delivers injected solutions to the chip surface creates ‘flow cells’. These are distinct volumes, each containing an individually monitored sensor surface. The experimentalist determines which flow cells receive solutions by selecting cells to define a ‘flow path’.

The surface within a flow cell is either a ‘reaction surface’, having some amount of functional immobilised ligand, or a ‘reference surface’, which has none. A flow path includes a reference surface and at least one reaction surface.

The simplest sensor chip is a glass slide with a thin gold film. Early sensor chips had a carboxymethyldextran (dextran) matrix bound to this sensor surface.^10^ Dextran is described in [5] (and in similar terms in [28]) as providing “… a flexible anchor for ligand immobilisation, allowing interactions to occur as in solution.” (For further details, see [29].)

More particularly, “The binding events [such as binding of analyte to immobilised ligand] occur in a threedimensional dextran matrix that extends approximately 100nm out from the sensor surface.” ([30]). We will call this region the ‘reaction volume’.

The monitoring of a flow cell results in a reported response that is updated in real time. This response may be influenced by changes in the mass present in a reaction volume. For example, the binding of analyte to immobilised ligand increases the mass in the probed volume. A corresponding increase in reported response indirectly shows the progress of the binding interaction.^11^ A graph of biosensor response against time is a ‘sensorgram’.

### 2.2 Interaction mechanisms

An experimentalist may expect that biomolecular interactions occurring within a reaction cell are described by some assumed system of chemical equations or ‘interaction mechanism’. For clarity, the mechanism should indicate details such as the valency of the immobilised ligand, and whether or not this ligand presents a homogeneous population of analyte binding sites.

One commonly used mechanism is the ‘simple bimolecular interaction’ (known by a variety of terms, such as the ‘one to one reaction’ [30]). This mechanism describes the reversible binding of analyte A and immobilised ligand B to form complex (AB), summarised by

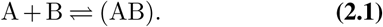

Another well known mechanism is the ‘two-state conformational change model’, also known as the ‘two-state model’ ([30]). Using the notation above, we may present this mechanism as

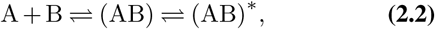

where (AB) and (AB)* are isomers.

### 2.3 Features of kinetic experiments

There are various types of assay — and types of experiments within these — designed for particular applications. We will consider data collected through a series of kinetic experiments in a ‘direct binding assay’ (DBA). The DBA is a fundamental form of assay as it uses an homogeneous analyte solution. The relative simplicity of a DBA allows us to define our experimental system in a relatively straightforward manner.^12^

A kinetic experiment has multiple phases, defined by the solution injected into the flow path. Prior to commencement of an experiment, a flow path is exposed to a running buffer solution that does not contain analyte. The experiment begins with an ‘association phase’. In a DBA this phase involves injection of a solution of known analyte concentration into the flow path for a specified period of time. The association phase is followed by a period known as the ‘dissociation phase’ in which buffer is returned to the flow path. An experiment may conclude with the injection of solutions through the flow path designed to dissociate complexes or clean the flow cells, followed by buffer. Only the part of the sensorgram relating to the association and dissociation phases provides data on the system kinetics.

Proceeding in this manner, the experimentalist injects solutions having a range of analyte concentrations, including a ‘blank’ having no analyte, obtaining a sensorgram for each. Ideally an experimentalist replicates the entire set of these experiments. Figure 2 presents a schematic version of a sensorgram obtained for a kinetic experiment as described that shows typical features of response.^13^

**Fig. 2.**
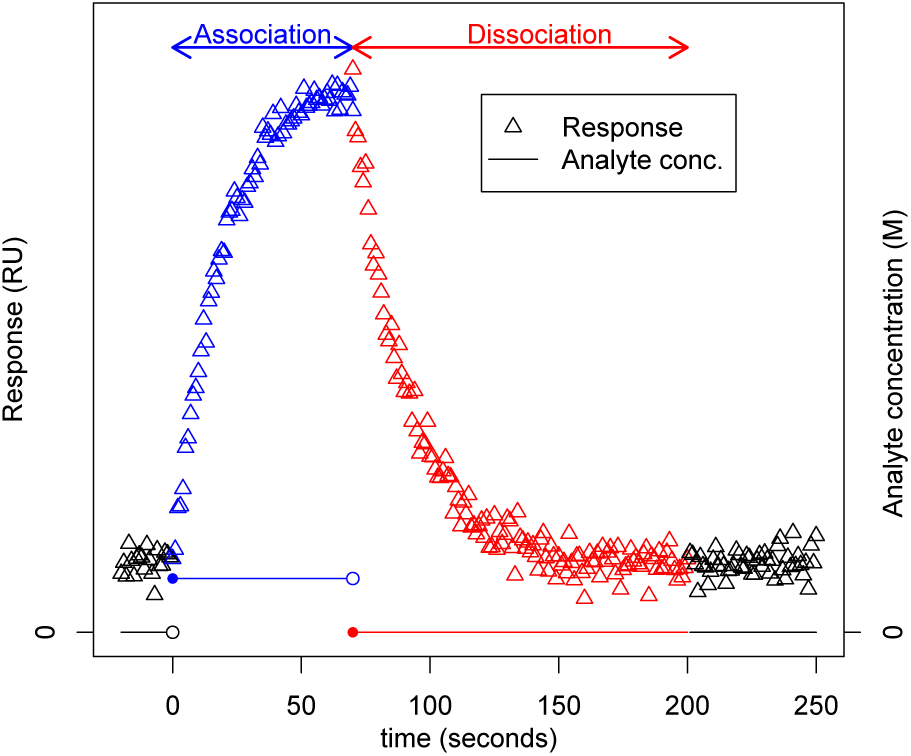
A simulated sensorgram for data subject to additive noise from a kinetic experiment in which the simple biomolecular interaction (Eq. (2.1)) applies. The time course of analyte concentration is superimposed on the graph to show changes in experimental conditions.

### 2.4 Components of experimental response

Recall that biosensor response (such as that having the general nature shown in Figure 2) has various components due to the multiple sources of mass in a reaction volume. Often the literature presents response expressions without making explicit mention of which components contribute to the modelled response.

Karlsson [30] provided a notable exception to this trend. There, “detector response” *R* was presented in terms of components due to “binding events and changes in buffer composition” and “baseline level” *R*_0_ via

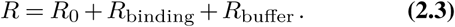

Later we will draw on Eq. (2.3) in forming a type of expression that is appropriate for our interest in specific contributors to biosensor response. For example, *R*_binding_ and *R*_0_ may be complicated due to the dissociation of immobilised ligand from the sensor surface, or being composed of multiple sub-components. We may make such dependencies explicit by employing a more general and more informative type of response expression. We will present this expression in Section 3.2.

The particular interest of Karlsson [30] was in the use of biosensor response expressions rather than their composition, as we are here. It suits our purposes to modify Eq. (2.3) to partition sources of response into a different set of components. This approach will allow us to more naturally define response components that we wish to remove from total experimental response when possible. If we can achieve this, we can readily determine whether we can simplify the response expression, addressing Q4.

In order to produce suitable terminology and notation for our discussion of response expressions, we will classify sources of experimental response and catalogue contributions to them. We consider four major components, each of which may be a sum of sub-components. Many of these contributors to response are described in detail in [5].

#### Response due to functional immobilised free ligand (*R*^FIL^)

If we know that the ligand is heterogeneous, then it is appropriate to allow a sub-component for each form of ligand. Heterogeneity may be due to chemically or structurally distinct forms of immobilised ligand. It is also caused when the immobilisation step produces multiple orientations of analyte binding sites which have varying accessibility to analyte.

#### Response due to complex (*R*^c^)

Our experimental system may have multiple forms of complex. Single complexes of these different forms may make unequal contributions to response.^14^ Further, the forms of complex may not have the same values for each of the rate constants that characterise the dynamics of the associated association and dissociation processes. Hence, it is appropriate to model the amounts of distinct complex forms separately. Accordingly, each distinct form of complex has its own sub-component in *R*^C^. Recalling Section 2.2, the model for the simple bimolecular interaction Eq. (2.1) has a single sub-component in *R*^C^ due to its one form of complex. The model for the two-state conformational change interaction Eq. (2.2) has two sub-components in *R*^C^ due to the two forms of complex.

#### Response due to non-interaction components (*R*^NI^)

This may be the sum of a variety of terms which include response due to:

- the sensor surface and matrix.
- injected solutions: contributions of analyte solutions and buffer to response may differ.
- sensor artefacts: examples include instrument drift and air bubbles following solution injection.
- non-specific binding: resulting from the binding of analyte to a site in the reaction volume other than the expected binding site(s) on the immobilised ligand, such as the matrix.

#### Response due to non-functional immobilised free ligand (*R*^NIL^)

Ligand immobilisation typically results in some amount of ligand that is unable to bind analyte. We model *R*^NIL^ by allowing one sub-component for each distinct form of ligand. It is appropriate to distinguish *R*^NIL^ from *R*^FIL^ as we expect the latter to change over time due to the interaction kinetics.

Informed by this discussion, we extend Eq. (2.3) and model biosensor response due to mass in a flow cell at time *t* as

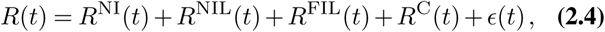

where ∊ represents random noise.

*Remark 2.1:* It is reasonable to expect that the form of ∊(*t*) in Eq. (2.4) will depend somewhat on the particular biosensor unit in use. It is not common for the literature to discuss experimental noise alongside a response model.

In simulation studies, authors may experiment with normally distributed additive noise in order to gauge its effect on parameter estimation from simulated data (see, for example, [32]). The Biacore™ Assay Handbook [33] provides guidance on expected features of residuals when a model appropriately captures the systematic features of data:

> Ideally, the residuals will scatter randomly around zero over a range that corresponds to the short-term noise in the detection system (approximately ± 1 to 2 RU). ([33, Appendix A1.4])

In deciding upon the form of noise for a simulation study, [34, page 178] performed an empirical study of BIAcore biosensor baseline response to inform their choice: “… an error with a normal distribution and a standard deviation of 0.584 RU …”, presumably with mean zero.

Eq. (2.4) is a flexible representation of response that is appropriate for various types of assays, interaction mechanisms, and experimental conditions. We will see in Section 3 that our approach makes model features explicit. This removes a barrier to investigating questions concerning model properties and appropriate data processing, as outlined in Q1-Q7.

### 2.5 Processing of biosensor response

Suppose we assume that biomolecular interactions occurring in a certain experimental series are suitably modelled by a particular interaction mechanism. We may use this mechanism to inform an expression for total experimental response based on Eq. (2.4) and our knowledge of the biosensor apparatus. We note that expressions for response in the literature generally do not include all of the components seen in Eq. (2.4). This may be because some components are constant, or otherwise do not provide useful information on the progress of interactions. As such, these ‘non-informative’ components may seem to be unwanted complications for any response expression, which may explain their frequent omission in the literature. However, actual biosensor data may contain such non-informative components. Hence, a model may not appropriately represent the data we collect. To remedy this issue, we need to appropriately process the data to remove non-informative components so that the result is suitable for parameter estimation using our assumed model (as we noted in Q1).

A typical means of processing data is by ‘reference subtraction’ (see, for example, [4]). This process is applied to all sensorgrams obtained from a reaction surface. One subtracts the sensorgram obtained from the reference surface under a given set of experimental conditions from the corresponding sensorgram obtained from a reaction surface.^15^ This subtraction is intended to remove *R*^NI^ from sensorgrams, which allows us to omit this component from a response expression. Such a model simplification may reduce the complexity of addressing Q4 subsequently. We note that appropriate data processing steps depend on features of the experimental system.^16^

In discussing the processing of response, we have only partially considered the practicalities of fitting a model to data. We still need to consider how to specify an explicit model. In order to achieve this, we must consider *R*^FIL^, *R*^NIL^, and *R*^C^ of Eq. (2.4) in more detail. Modelling these components requires an understanding of the relationship between mass near a sensor surface and the associated response.

We consider this matter as part of a review of response expressions in the literature. We undertake this review to demonstrate the diversity of models employed, and how they tend to have implicit features. These features complicate the understanding of relationships between interactions and response, and are also responsible for Q1-Q7. We intend that our review will demonstrate the value of developing an unambiguous system for representing biosensor response.

We begin our discussion of response expressions by presenting relationships between biosensor response and amounts of interactants seen in the literature. This aids a comparison of the properties of these alternative expressions, and informs the response modelling framework we propose subsequently.

### 2.6 Relationships between biosensor response and experimental quantities

#### 2.6.1 Response due to surface mass concentration of a species

The biosensor response due to a particular source of mass near a sensor surface is directly proportional to its surface mass concentration ([1]). Let us illustrate this relationship by considering some immobilised species X, and the response due to this, say *R*_*X*_ in response units (RU). To draw on the notation of [1], we relate *R*_*X*_ to the surface mass concentration of X, say Γ_*X*_, having units of g.m^−2^, by

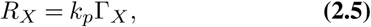

where *k*_*p*_ is a constant of proportionality with imputed units of RU.m^2^.g^−1^.

#### 2.6.2 Response relating to molar concentration of a species

Statements relating surface mass concentration to response (as in Eq. (2.5)) are pervasive in the literature. By comparison, relationships between molar concentration and response are seen relatively rarely. From these we note two key properties (P1 and P2 below) that we will find useful subsequently. (P1) O’Shannessy and Winzor [35] asserted that the response component due to complex is directly proportional to the molar concentration of that complex.

Karlsson [30] considered molar concentration of analyte explicitly in his consideration of Surface Competition Assays (SCAs).^17^ As the two analytes present had unequal molecular masses, Karlsson’s model had to account for their differential effect on response. To explain this, consider some analyte X, and following Karlsson’s notation, let *n* denote the refractive index of a reaction volume. Changes to *n* due to binding of analyte to immobilised ligand depend on the analyte concentration, *c*_*X*_ (g ⋅ ml^−1^, a mass in a volume, defined in Eq. (1.3)) and the refractive index increment, *dn/dc*_*X*_, through

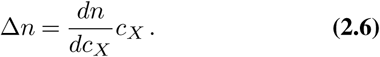

This expression led to the description of the response due to binding of analyte, *R*_binding_ of Eq. (2.3). Karlsson also asserted: (P2) “When analytes with similar refractive index increments are considered, and when the concentration of the analyte is expressed in molar terms, this means that *Δn* and therefore also *R*_binding_ will be proportional to the molecular weight of the analyte.” ([30])

We will now demonstrate when P1 and P2 are compatible. Let us consider some species X for which Eq. (2.6) holds, and recall the notation of Section 1.1. We may use Eq. (2.6) to relate the response component attributable to X (*R*_*X*_) to *M*_*X*_ and the molar concentration of X, [X].

Rearranging Eq. (1.1) into *m_X_ = N_X_ M*_*X*_, we rewrite Eq. (2.6) as

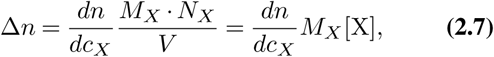

for [X] as defined in Eq. (1.2).

Following P2, *R*_*X*_ is proportional to Δ*n*. We use Eq. (2.7) to convert this into an equality with constant of proportionality *η*:

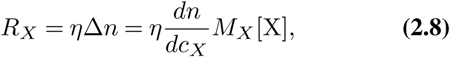

which is consistent with P1.

*Remark 2.2:* Suppose we use Eq. (2.8) to compare the individual responses due to two species that have the same molar concentration and refractive index increment but have differing molar masses. This situation makes it apparent that response due to a species is proportional to the molecular mass of that species.

We will now rewrite Eq. (2.8) to create a simplified relationship that we will employ subsequently.

*Assumption 2.1* (Response due to a bound species) Suppose species X is either free immobilised ligand or analyte-immobilised ligand complex. Further suppose *M_X_* exceeds the biosensor’s detection limit. In this case, [X] (units M) and *R*_*X*_ (RU) are related by

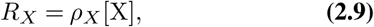

where *ρ*_*X*_ has units RU. M^−1^ Comparison of Eq. (2.9) and Eq. (2.8) shows that *ρ*_*X*_ depends on two properties of X: its refractive index increment, and its molecular mass.

We will see in Section 3 that Eq. (2.9) contributes to the transparency of our modelling framework.

To the best of our knowledge, the literature has not compared the relationship between the molar concentration of a surface-bound species and resultant response (as in Eq. (2.9)) with the relationship between surface mass concentration and response (as in Eq. (2.5)). Below we establish conditions under which the two expressions are equivalent. This may assist the reader in comparing some model forms seen in the literature.

#### 2.6.3 Reconciling relationships between bound species and associated response

For some immobilised (or bound) species X, let us consider the relationship between [X] and Γ_*X*_.

Consider a reaction volume of *V* (dm^3^). For *s* (m^2^) representing the sensor surface area,

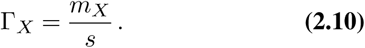

Employing Eq. (1.1) and Eq. (1.2) we derive

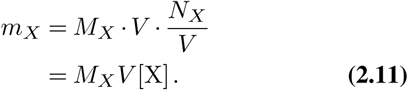

Substituting Eq. (2.11) into Eq. (2.10) yields

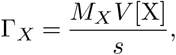

and hence from Eq. (2.5)

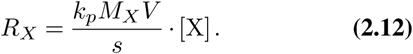

As *M*_*X*_ is constant, if (*k_p_. V*)/*s* is also constant, then so is the coefficient of [X] in Eq. (2.12). Thus, Eq. (2.12) has the same form as Eq. (2.9). This shows when Eq. (2.5) and Eq. (2.9) are consistent. Further, Eq. (2.12) demonstrates that the component of response due to species X in the reaction volume is directly proportional to both *M*_*X*_ and [X], as seen in Eq. (2.8). These observations support the claims made in P2 and P1 respectively.

Relationships such as those presented in this section have found use in the various representations of biosensor response seen in the literature. We now proceed to give an overview of the major types of these. This summary will demonstrate some limitations that we will seek to avoid in our formulation of models for response data in Section 3.

### 2.7 Representations of biosensor response in the literature

Some papers present an explicit expression for response as a function of time ([36, 37]). The complexity of such expressions, alongside their tendency to have unexplained features, tends to make answering Q1 and Q3 unnecessarily difficult. This lack of detail also complicates attempts to address questions amongst Q4-Q7.

Other forms of models for response components may be easier to compare with each other. However, prior to such comparisons, we may need to understand the meaning of certain terms (an instance of Q2). We may not readily gain this understanding due to the manner in which response models are presented. In order to demonstrate this matter, we will consider examples of two broad types of response expressions.

#### 2.7.1 Models relating rate equations to response

A paper may provide an ODE, or a system of these, describing the rates of change of variables. Each of these variables represents some amount of a chemical species. We may find the meaning of these variables unclear when terms are not defined (examples include [34, 38, 39]). At other times, such as when a variable is described as a ‘concentration’, the reader can infer the meaning of the term. For example, [40] presented a rate equation for “the concentration of complex [AB]” assuming that the simple biomolecular interaction (Eq. (2.1)) was appropriate. The equation featured terms for [AB], and also free analyte A and immobilised ligand B, represented by [A] and [B], respectively. For [A] given units M and the forward rate constant given units of M s^−1^, we infer that the rate equation describes a reaction occurring in a volume (as cited in Section 2.1). The combined effect of these cues is to suggest that [A] and [B] each represent a molar concentration.

Karlsson *et al.* [40] proceeded to relate [AB] to the response due to bound analyte by citing a relationship akin to Eq. (2.5). This type of relationship is often cited when papers convert a rate equation system into a response ODE system.^18^ However, we may have some reservations about the use of the relationship in [40] given that the original equation was in terms of molar concentration variables, rather than mass concentration variables as employed in Eq. (2.5). Following our discussion in Section 2.6.3, we see that we can express a response component as proportional to either mass concentration or molar concentration. However, the constant of proportionality used in response simulation (or sought in parameter estimation) will depend on which of the two relationships we employ.

Some papers provide little information on how they convert a rate equation to a response equation. A paper may justify the conversion via a statement such as

> The signal observed, *R,* is proportional to the formation of AB complexes at the surface… ([39] describing response resulting from the simple biomolecular interaction, Eq. (2.1))

There are various instances in which some unspecified relationship is used to translate rate equations into ODEs for response components.^19^ In such cases it appears that the latter is obtained from the former by directly replacing each variable representing an amount of immobilised or bound species by some corresponding response variable.

We see a variant of this issue when a paper that begins with an ODE system in concentrations of free binding sites proceeds to present an expression in RU. This expression may be a solution of the original ODE system transformed into response through the use of coefficients having units of RU (see, for example, [42]). This may conceal how we may derive a response equation from the rate equations.

A similar matter arose in [26]. There, rate equations were given for two immobilised species: AB, a ligand which is an antigen-antibody complex, and CAB, a complex between the ligand and chemical species Fab. These species contributed to (seemingly processed) response at time *t, R_t_,* which had the form^20^

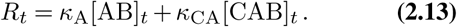

As noted for other models, the variables (here [AB]_*t*_ and [CAB]_*t*_) were not defined.

The paper advised that terms such as κ_A_ and κ_CA_

> … are used to account for the difference in mass between the antigen alone and the Fab-antigen complex.

This description indicates that [AB]_*t*_ and [CAB]_*t*_ themselves are not mass terms. Instead, Eq. (2.13) used the counterparts of κ_A_ and κ_CA_ to associate the effect of molar mass with [AB] and [CAB], producing a response component for each species. Comparing Eq. (2.13) and Eq. (2.9), we may reason that [AB]_*t*_ and [CAB]_*t*_ are molar concentration terms.

We note that [26], by differentiating between chemical species that make different contributions to response (via an expression like Eq. (2.13)), employed a more general modelling approach than others reviewed in this section. Significantly, [26] demonstrated the importance of taking account of the differing contributions to response when replacing each distinct concentration term with a corresponding response term, in order to ensure that we arrive at a correct response expression. From this we surmise that it is certainly not appropriate to automatically replace the symbols representing concentration variables in rate equations by others for response variables in all situations. We note that the frequent application of this practice to simpler models in the literature conceals the importance of the modelling approach of [26] for more complex models.

*Remark 2.3:* The parameters κ_A_ and κ_CA_ in Eq. (2.13) are effectively constants of proportionality between some (molar) concentration and the response due to it, but they are not formally defined. This creates some uncertainly over the meaning of the modelled response in Eq. (2.13).

We cannot be certain if the intent of Eq. (2.13) is to present an expression for *R*^FIL^ + *R*^C^ (as occurs in Eq. (2.4)), or something else. From the notation, we may surmise that κ_A_ and κ_CA_ only model the response due to bound ligand, and do not include contributions from the mass of immobilised ligand. In this case, we may expect that the model represents response processed to yield *R*^C^ only.

Understanding the modelled response is important as it has implications for the meaning of parameters. Suppose that Eq. (2.13) does represent *R*^*C*^. Then, we must take care over how we use a relationship such as Eq. (2.9) to relate a molar concentration to a response component. We must modify it appropriately so that it depends on the molar mass of A or CA, rather than AB or CAB, respectively.

Aside from the matters above, there is another benefit to clearly defining model terms. The accompanying certainty allows us to relate parameters such as κ_A_ and κ_CA_ to physical quantities. This will assist us in parameter estimation by guiding us towards reasonable initial guesses for the value of these parameters. Having these starting values may result in a parameter estimation algorithm converging to an estimate of the parameter vector in fewer iterations than otherwise.

Certain authors have provided specific guidance to assist model interpretation. For example, Haake *et al.* [41] indicated that their response equation modelled response due to bound analyte (there, “ligate”). However, other works are unclear as to whether response is due to bound analyte or analyte-immobilised ligand complex ([34, 38, 39]).

#### 2.7.2 Models for response only

Models for response components are often presented as a first-order ODE (systems of these are much less common) for the response due to a particular chemical species. Such an equation may have the appearance of a chemical rate equation due to the notation employed. At times, variables are described, such as in [43]:

> The amount of free receptor sites on the surface ([B]) and the response attributed to the IL-2-receptor complex ([AB]) were modeled in response units (RU).

Also, we can infer that analyte concentration [A] in [43] represents a molar concentration from the description of experiments. In describing the simulations which used their ODE model for the two-state conformational change interaction Eq. (2.2), [44] indicated that injected analyte (there, “ligate”) and “total matrix site concentration” (related to the amount of immobilised ligand) had units appropriate for molar concentrations.

At other times terminology is less clear or units are not specified (recall Q2). In [14] “concentration” was used to describe both analyte concentration [A] and the concentration of free receptor sites [B]. Further, there may not be any indication of which variables contribute to response, or the manner in which they do this. (See, for example, [14, 30, 44].) In other works, an ODE system for response components due to forms of complex may introduce a term to account for differences between them. However, the authors may not relate this term to anything concrete that would guide the composition of a response model for a similar interaction mechanism. (For example, see the “normalising factor” employed in [45].)

Such instances of incomplete model description are an obstacle to model interpretation. The uncertainty to arise from this outcome may lead to questions from amongst those in Q1-Q7 listed in Section 1. Uncertainty around the meaning of models is exacerbated by the general aversion of the literature to modelling the multi-component nature of experimental response. Model interpretation is assisted when components are made explicit, as occurred in [40], which we summarised in Section 2.7.1. We will draw on and extend this example in Section 3.2. This extension will lead to a means of specifying models for biosensor response more precisely. Further, it will enable the testing of models for certain properties. We will consider this task next when we review the state of the art of methods for testing models of biosensor response for global *a priori* identifiability.

### 2.8 Testing models of biosensor response for global a priori identifiability

Testing mathematical models of biological systems for the property of global *a priori* identifiability began with [46].^21^ The authors considered a (linear, time-invariant) model for the movement of radioactive tracers between compartments of an organism.^22^ Since that time, researchers have developed methods for analysing a wide range of model classes, and applied these methods to models from various fields.

Testing models of biosensor data for global *a priori* identifiability poses certain challenges. The abrupt change in experimental conditions that delineates the phases of a kinetic experiment (as shown in Figure 2) led to the classification of some models as a type of linear switching system (LSS). These systems are not suited to the standard methods of analysis designed for linear time-invariant or nonlinear models. However, [7-9] proposed methods that were sufficient to test a form of model for the simple bimolecular interaction (Eq. (2.1)) and the two-state conformational change interaction (Eq. (2.2)). Recent developments suggest that there is scope for further progress in this area ([10]). These developments will assist the analysis of those models which can be presented as an LSS. Such a model form follows readily from the modelling framework we propose here.

Due to the variety of representations of models for biosensor data, the early efforts to test linear switching systems representing biosensor data for global *a priori* identifiability in [7-9] sought to make the task as simple as possible. This was achieved by considering simplified forms of the models, and omitting model derivations. As a result, [7-9] addressed Q5, but not Q4 and Q7. A consideration of the simple bimolecular model in [10] showed that the answer to Q7 is ‘yes’. We strongly suspect that asking this question of other models will yield useful insights into their properties.

We place ourselves in a position where we may begin to address Q1-Q7 by formalising aspects of biosensor response models. We propose a means of achieving this in the next section.

## 3 Towards an unambiguous description of biosensor data

Recall from Section 2.7.1 the notion of a parametric mathematical model that relates biosensor response to amounts of chemical species. We propose that this idea points towards an effective way to clearly present a model. More particularly, we propose that clarity is served by using some assumed interaction mechanism and the experimental conditions to inform a model of two parts.

The first, a ‘kinetic model’, is a description of the amounts of biochemical species over time. Recall that species interactions occur in a three-dimensional volume. As such, it is natural for a kinetic model to employ rate equations (ODEs) for molar concentrations of interacting species. The second, a ‘response model’, is an expression for biosensor response that has components due to various sources of mass in a flow cell. It is based on the general model presented in Eq. (2.4). In order to relate the response model to the kinetic model, we require a relationship between response due to an interacting species and its molar concentration, as in Eq. (2.9).

Henceforth we shall term the combination of a kinetic model and a response model a ‘kinetic-response model’. In mathematical terms we classify this as a state-space system. This classification is advantageous as there are various techniques available for the analysis of such systems, including methods of testing them for global *a priori* identifiability. More generally, in earlier studies ([7-10]) we found that expressing biosensor response as a kinetic-response model assisted us in addressing aspects of Q1-Q7 subsequently.

We will not consider a particular interaction in detail here as the variety of models seen for even the least complex interactions warrants a separate, detailed review.^23^ Instead, we present a general form for each of a kinetic model and a response model. We then show how these two models are related. We illustrate the form of a kinetic-response model with an example. We conclude the section by demonstrating how to reduce this initial model to a simpler model when data is collected under suitable experimental conditions.

### 3.1 A general kinetic model

A kinetic model concisely presents the rate equations for certain species. We limit our interest to forms of free immobilised ligand and complex.

Let 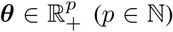 denote the vector of all parameters that will appear in a kinetic-response model. This vector includes rate constants, unknown initial molar concentrations of interacting species, and other parameters that relate interactant concentrations to components of response. Molar concentrations of interacting species in the reaction volume at time *t* are represented by the state vector at time *t,* 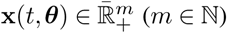. We may write the rate equations for all species in x as a first-order ODE system of the form

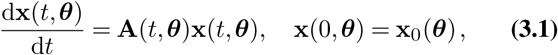

where elements of 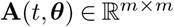 are functions of the rate constants in *θ* and the time course of analyte concentration. Matrix A in Eq. (3.1) is “compartmental” (see, for example, [49]) which ensures that mass is conserved in the model. This feature prevents the model from displaying unphysical behaviour, such as by predicting negative concentrations. The size and elements of A are determined by the particular interaction mechanism assumed.

### 3.2 A general response model

Recall the response components as described in Section 2.4, and combined in the general response equation Eq. (2.4). We modify the latter to show parameter dependence:

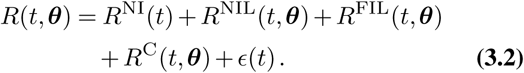

When considering a particular interaction mechanism, we would produce a suitable kinetic model having the form Eq. (3.1). The particularities of this kinetic model would direct us in adding detail to the general response model Eq. (3.2), resulting in an appropriately informative response model.

When there are multiple forms of functional immobilised free ligand or complex, we will gain further clarity by representing those (meaningful) components of response, R^FIL^ or R^C^ in Eq. (3.2), by a sum of sub-components as is appropriate.

We may use Eq. (2.9) to represent that part of response Eq. (3.2) due to forms of complex and immobilised ligand as a function of x by

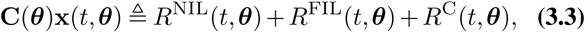

where 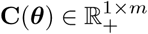. We omit *R*^NIL^ from Eq. (3.3) if the amount of non-functional ligand is constant, and regard it as a non-informative component of Eq. (3.2) instead.

Next we shall illustrate the clarity afforded by a kinetic-response model. In the interests of presenting a simple example, we will consider a case in which we can neglect complications due to non-ideal experimental conditions.

### 3.3 A kinetic-response model example

#### 3.3.1 Assumptions

Suppose we consider a kinetic experiment in a DBA where total response is described by Eq. (3.2), and:

- the simple bimolecular interaction mechanism Eq. (2.1) is appropriate,
- reference subtraction allows us to process any sensorgram to remove R^NI^(*t*, θ),
- the immobilised ligand:
  − is completely functional, that is, *R*^NIL^(*t*, *θ*) ≡ 0,
  − is homogeneous, hence *R*^FIL^(*t*, *θ*) has one component,
  − cannot dissociate from the surface, and
  − does not have any analyte bound at the start of an experiment.

Under these assumptions, we arrive at an expression for processed response at time *t*:

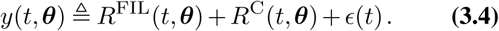

In Eq. (3.4), *R*^FIL^(*t*, *θ*) + *R*^C^(*t*, *θ*) is that part of Eq. (3.2) that relates to the progress of the interactions. All terms in Eq. (3.4) are time-varying. (For example, we expect R^FIL^ to decrease as the association phase proceeds whilst R^C^ increases.)

#### 3.3.2 The kinetic model

We assume that the molar concentration of free analyte A is constant in the association phase of an experiment. Following Eq. (3.1), the kinetic model (relating the concentrations of immobilised interacting species) has the form

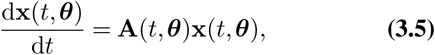

where

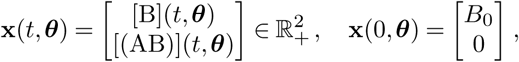

and parameter *B*_0_ > 0 has units of M.

#### 3.3.3 The response model

Following Eq. (3.4), we use Eq. (3.3) with *R*^*NIL*^ ≡ 0 to relate the response model to x from Eq. (3.5), yielding

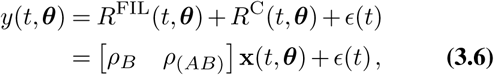

where 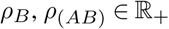 draw on the convention of Eq. (2.9).

#### 3.3.4 The kinetic-response model

By consolidating Eq. (3.5) and Eq. (3.6) into one state-space system we obtain the kinetic-response model

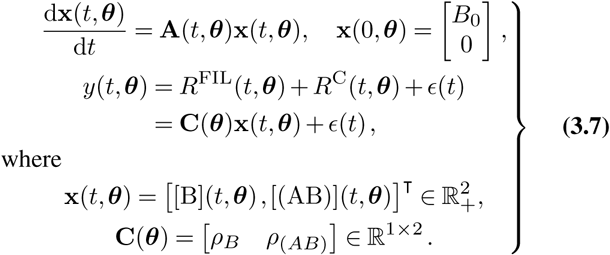

We will see in Section 3.4 that our approach to defining a kinetic-response model may allow subsequent simplification. We will demonstrate this simplification by using Eq. (3.7) as an example.

### 3.4 Further simplification of models and associated processing of data

Consider the kinetic-response model Eq. (3.7). We can use our knowledge of the biosensor to express certain model features in more fundamental terms. We will see shortly that this approach enables us to simplify our kinetic-response model.

To begin the process of model simplification, we consider the parameters *ρ*_*B*_ and *ρ*(*AB*). By not merely regarding these as constants of proportionality, ultimately we can show a relationship between them.

*Remark 3.1:* In the interests of simplicity, let us assume that both B and (AB) have the same constant refractive index increment, say *k.* As a result, following Eq. (2.8) and Eq. (2.9) and employing a constant 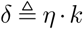 we note that

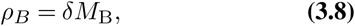

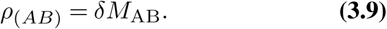

That is, due to its greater molecular mass, the contribution of one (AB) to response is greater than that made by one B.

Our model simplification is also assisted by considering the initial response predicted by Eq. (3.7).

*Remark 3.2:* Let us define the time immediately before the injection of analyte in an experiment as *t* = 0. From the response expression in Eq. (3.7),

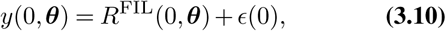

and from Eq. (3.5) and Eq. (3.6),

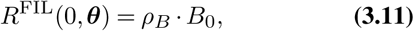

showing that we can express this response component purely as a product of parameters.

We may use Eq. (3.9) and Eq. (3.8) to write

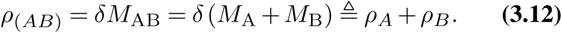

Hence, we can rewrite the response in Eq. (3.7) as

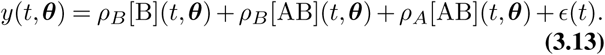

We note that *ρ*_*B*_ [B](*t*, *θ*) + *ρ*_*B*_ [AB](*t*, *θ*) in Eq. (3.13) is the total contribution to response by B from its free and bound forms, that is, *R*^FIL^ (0,*θ*) as in Remark 3.2, Eq. (3.11). Under our modelling assumptions, this is a constant, positive value. Hence, by subtracting Eq. (3.11) from Eq. (3.13) we obtain a processed form of response:

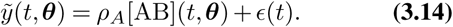

We interpret Eq. (3.14) as the response due to bound analyte subject to noise.

This derivation produces a response model Eq. (3.14) that has one fewer parameter than Eq. (3.6). Further, we have demonstrated a benefit of our formulation of a kinetic-response model. We can see how to reduce our original response model Eq. (3.6) to an expression that is consistent with the form employed by [30], as illustrated by Eq. (2.3). That is, we have shown how our kinetic-response model formulation assists us with addressing Q1, Q3 and Q4. From this point we can proceed to consider questions such as Q7.

A process of simplification that is similar to that shown above may apply to other cases where the response model has a greater number of components. For example, we could apply very similar arguments in simplifying the response model Eq. (3.3) for the simple biomolecular interaction Eq. (2.1) having constant and positive *R*^NIL^ (*t*, *θ*).

## 4 Discussion and Conclusions

The literature shows a variety of approaches to writing response models, as we saw in Section 2.7. These approaches tend to omit important features that would make their meaning clear. This lack of clarity inhibits scientific communication, and makes it unnecessarily difficult to test alternative model forms for equivalence.

An incompletely specified model also impedes our ability to inspect model properties. Of these, it is particularly useful to test an assumed model for the property of global *a priori* identifiability. Such a test can foresee the maximum amount of information we can expect to obtain from a planned series of experiments. For example, the test may allow us to anticipate which experiments cannot lead to a unique estimate for each parameter. Further, the result of a test can highlight specific deficiencies in the combination of the assumed model and experimental design, and suggest remedies for these. This knowledge can guide us towards a reformulation of our model or an alternative experimental design. As a result, we can arrive at a planned study that has at least the potential to provide useful information on our system, whereas the original study almost certainly could not.

In this paper we have outlined a framework for specifying response models in an unambiguous manner. Towards this, we have formulated a general biosensor response model Eq. (3.2). We may use this to write an appropriate response model for a biomolecular interaction system regardless of the assumed interaction mechanism.

Our framework employed a model of two parts. The ‘kinetic model’ is a system of first-order ODEs for the rates of change of the molar concentrations of the immobilised species that participate in interactions. We connected this to a ‘response model’ that showed unambiguously how the species in the kinetic model contributed to biosensor response. We termed the combination of the kinetic model and the response model a ‘kinetic-response model’, and noted that it had the form of a state-space system.

Our previous investigations of the kinetic-response models associated with particular interaction mechanisms have demonstrated the benefits of this model form ([10]). Significantly, the form is amenable to testing for global *a priori* identifiability. Also, compared to other types of models, it is much easier to survey the features of a kinetic-response model. This can lead to model simplifications in certain experimental situations. This is useful as preliminary studies have shown that some original kinetic-response model may lack the property of global *a priori* identifiability, whereas a simplified model derived from it has the property [(10, Chapter 5]). That is, it may be at least possible to uniquely estimate all rate constants of a reformulated kinetic-response model when this is almost certainly impossible for the original model.

Our formalism for specifying response in terms of components is general enough to suit a diversity of experimentally observed situations. These include the cases where immobilised ligand is heterogeneous, or has a portion that is non-functional, or dissociates from the sensor surface over time. Our template is not only appropriate for modelling the response obtained from a DBA. It is also suited to modelling response from those assays that employ a heterogeneous analyte solution, such as the SCA. When relating a response model as in Eq. (3.2) to the kinetic model (having general form Eq. (3.1)), we expect that having a relationship between parameters and physical quantities will assist parameter estimation (see Remark 2.3).

Progress towards defining a kinetic-response model for a particular interaction mechanism will require a suitable kinetic model. The diversity of representations we noted for response models in Section 2.7 is also seen in the description of biomolecular interaction dynamics associated with interaction mechanisms. Through work in progress, we aim to reconcile alternative models associated with the simple bimolecular model ([47]) and the two-state conformational change model ([48]). This will enable us to completely specify an appropriate kinetic-response model for each.

There are a number of flow-cell optical biosensor interaction models which are more complex than the two referenced in Section 2.2. (See, for example, [14, 15, 30, 42, 45]). Our scrutiny of simpler interactions here will provide a template to guide the complete specification of models for more sophisticated interactions. Having achieved this, we may also be able to test these models for global *a priori* identifiability using the approaches of [9, 10].

We note that ‘equilibrium experiments’ are performed to provide data for use in estimating equilibrium constants. Adapting the terminology introduced earlier, we may propose that such data is modelled by an ‘equilibrium-response’ model. The analysis of such models is simpler than that for kinetic-response models associated with kinetic experiments. Accordingly, our discussion here will readily translate to a means of specifying equilibrium-response models that is suitably informative.

## Acknowledgements

The LATEX template for this manuscript was adapted from one made publicly available by Ricardo Henriques [50].

The author is grateful for support for the original research from Professor Anthony Guttmann through a postgraduate scholarship from the Australian Research Council Centre of Excellence for Mathematics and Statistics of Complex Systems at the School of Mathematics and Statistics, University of Melbourne. Subsequent refinements and the writing of this paper were made possible by a Faculty of Science Writing-Up Scholarship supported by the David Lachlan Hay Memorial Fund (University of Melbourne) and a Lift-Off Fellowship from the Australian Mathematical Society.

## Footnotes

1 Due to their means of detecting biomolecular interactions, these are also known as surface plasmon resonance (SPR) biosensors. For an introduction, see [1].

2 See the bibliography of [3] which has an extensive section for references relating to Biacore™/GE Health biosensors.

3 We show this mechanism in Section 2.2. This mechanism is commonly chosen (at least initially) as a suitable model to fit to data when we assume that certain conditions hold. A fundamental assumption is that the interaction has a 1:1 stoichiometry.

4 The test of a model for identifiability proceeds subject to certain assumptions. One of these is that we consider an infinite, error-free output trace. The assumption is reasonable; if we find multiple parameter vectors which can produce such an idealised trace, this non-uniqueness is also very likely to occur when data is limited in extent and subject to error

5 We propose that the easy-to-use nature of BIAevaluation software coupled with an algorithmic approach to data processing that is appropriate for some — but not all — models has concealed the benefits of the model analysis we advocate for here.

6 One situation where this occurs is where ligand immobilisation is achieved by an ‘antibody affinity capture approach’, see, for example, [21]. In such a case, quantifying kinetics with the (surface) ‘decay model’, discussed in [26], may be appropriate.

7 Wang *et al.* [17] considered fitting a model for an interaction they termed the “1:1 binding model” (likely equivalent to Eq. (2.1)) to data. They noted that: “Comments on more complex binding models can be input through Biacore or other evaluation software.”

8 We appreciate that the experienced biosensor practitioner will be familiar with the background information we present. However, we note that these preliminary details are distributed through the literature. We consider that collecting the necessary information together in one place and formalising it where necessary will assist the reader in appreciating this paper’s novel contribution.

9 Much more widely cited is a relationship between response and mass concentration of a species.

10 There are a variety of sensor surfaces to facilitate specific applications; see [27] for a review, or the summary [21, Table 1].

11 Assuming that the mass increase is above the instrument’s lower detection limit; see [5].

12 We direct the reader interested in a description of assay types to [31]. We note that both the simple biomolecular in eraction and the two-state conformational change interaction may be appropriate descriptions of biomolecular interactions occurring in a DBA.

13 We note that real data may be subject to experimental effects, such as spikes in response caused by a change from one injected solution to another. We consider these further in Section 2.4.

14 Response due to a species depends on its effect on refractive index near the sensor surface. Thus, the contribution due to a form of complex depends on its refractive index increment and molecular weight; see [30] for details. We will use these properties in Section 2.6.2.

15 Relevant experimental conditions include the injected analyte concentration, response due to immobilised ligand on the reaction surface, and solution flow rate.

16 Recall our mention in Section 1 of experiments where immobilised lig- and dissociates from the sensor surface. We intend to address this case when it is appropriately modelled as in [26] in a separate study.

17 A SCA employs a heterogeneous analyte solution in the association phase of an experiment. The analyte species are assumed to bind immobilised ligand but not each other. See [31] for further details and assay types

18 See, for example, [41], in which we interpret “surface concentration” as mass concentration.

19 See, for example, [38] which considered the “1:1 Langmuirian interaction model”, likely equivalent to the simple bimolecular interaction model.

20 The original notation of [26] had R_A_ and R_CA_ as coefficients of [AB]_*t*_ and [CAB]_*t*_ respectively. We replaced this by the notation in Eq. (2.13) to prevent possibly confusion with response components in Section 2.4.

21 The authors termed the property “structural identifiability”. There are other terms in the systems theory literature, which are not always equivalent.

22 Although the organism was not specified, models from this broad class have found use in the study of tracer movement in various species, including human subjects.

23 These are in progress for the simple bimolecular interaction Eq. (2.1) ([47]) and the two-state conformational change interaction Eq. (2.2) ([48]).

